# Ectopic expression of the sensor CovS cytosolic domain confers phosphatase activity and full virulence to the M1T1 CovSY39H attenuated variant of group A *Streptococcus*

**DOI:** 10.1101/2025.07.30.667676

**Authors:** Abhinay Sharma, Aparna Anand, Miriam Ravins, Usha Kantiwal, Emanuel Hanski

## Abstract

Group A *Streptococcus* (GAS) causes a wide variety of diseases ranging from mild, noninvasive, such as pharyngitis and impetigo, to life-threatening infections, such as necrotizing fasciitis (NF) and streptococcal toxic shock syndrome (STSS). The two-component CovR/S system, comprising the sensor kinase CovS and transcription factor CovR, is a central regulator of GAS virulence. An attenuated pharyngeal colonizing variant (S126) possessing a single-nucleotide polymorphism (SNP) in CovS (Y39H) was recovered in France from a member of a family in which another individual developed NF and STSS caused by the M1T1 WT strain (S119). We employed transcriptome analyses (RNA-seq), quantitative determinations of CovR phosphorylation, measurements of virulence factor activity, and a murine model of human NF to demonstrate that CovS of strain S126 almost lost its entire phosphatase activity but retained its kinase and phosphotransfer activities. Moreover, we reversed its attenuated phenotype by ectopically expressing the cytosolic domain of wild-type CovS. Culturing the corresponding strain S126*cov*S-3’ in a chemically defined medium (CDM) supplemented with asparagine (Asn), conditions that produce an excess of cytosolic ADP over ATP, stimulated the ectopically expressed phosphatase activity. Consequently, this led to dephosphorylation of CovR∼P and increased the expression of virulence factors. Most importantly, S126*cov*S-3’ reverted to the wild-type phenotype of S119 in the mouse model of human GAS NF. Our study provides a new mechanistic tool that enables the manipulation of CovS phosphatase activity both *in vitro and in vivo*.

**IMPORTANCE:** The shift from high to low virulence and back in GAS is critical for understanding its pathogenesis and developing new treatments to control GAS infections. The two-component CovR/S system, comprising the sensor kinase/phosphatase CovS and the transcription regulator CovR, regulates the degree of GAS virulence. In the M1T1 serotype, *cov*R/S mutations are typically associated with a loss of CovR/S function, leading to hypervirulent phenotypes. However, an attenuated pharyngeal-colonizing variant possessing a single-nucleotide polymorphism (SNP) in CovS (Y39H strain S126) was isolated in France. Here, we demonstrate that S126 is attenuated because the mutation in CovS inhibits its phosphatase activity but preserves its kinase and phosphotransfer activities. By expressing the cytosolic domain of WT CovS in the S126 background, we endowed the resulting strain, S126*cov*S-3’, with phosphatase activity, which was further stimulated by ADP when formed in chemically defined medium (CDM) supplemented with asparagine (Asn). Most importantly, S126*cov*S-3’ switched to full virulence, similar to that of S119 in a mouse model of human GAS NF. These findings underscore the importance of comprehensive analyses of disease-related *cov*R/S mutants in understanding the virulence and persistence of GAS.

## INTRODUCTION

*Streptococcus pyogenes*, also known as Group A *Streptococcus* (GAS), is an extracellular, human-adapted pathogen recognized as one of the top ten causes of infectious mortality (1). GAS causes many human diseases, ranging from pharyngitis and impetigo to life-threatening infections such as bacteremia, necrotizing fasciitis (NF), and streptococcal toxic shock syndrome (STSS) (1, 2). Repeated and untreated GAS pharyngeal infections can result in immune response-mediated pathologies that can progress to chronic autoimmune conditions, such as rheumatic heart disease (RHD) (1). An estimated 616 million infections and more than 500,000 deaths per year are attributed to GAS infections (3).

Among the more than 200 known serotypes, M1 is the most frequently identified serotype to be associated with invasive diseases (4, 5). A hyperinvasive M1T1 clone was first detected in the mid-1980s in the United States and has disseminated worldwide, causing NF and STSS (6, 7). Since 2008, a subclone of M1T1 (M1UK) has emerged, significantly contributing to scarlet fever outbreaks in the United Kingdom and, more recently, causing a surge in invasive infections worldwide (8–14).

Control of virulence (CovR/S) or capsule synthesis regulator (CsrR/S) is the best-characterized two-component system (TCS), which is the primary regulator of GAS virulence (15, 16). It comprises a sensor kinase, CovS, and the transcription regulator CovR. It regulates up to 15% of the GAS genome (15, 17, 18) and coordinates the response to stress conditions, such as limited nutrient availability and host-pathogen interactions (19–23). When CovR is phosphorylated, it binds to the promoters of numerous virulence-associated genes, blocking their transcription, whereas dephosphorylation leads to their expression (24).

CovS shares domain organization and structural similarities with the EnvZ and PhoQ sensors of a two-component signal transduction family, which possess histidine kinase (HK) and phosphatase activities (25, 26). ATP is a substrate for HK, whereas ADP stimulates its phosphatase activity (27–31). Cathelicidin host-defense peptide (LL-37) binds to the surface-exposed CovS domain, increasing its phosphatase activity and thus enhancing the transcription of virulence factors (2, 22, 32, 33). Mutations in *cov*R/S can occur spontaneously during M1T1 infections and typically result in hyper-virulence due to the loss of all CovR/S functionality (5, 34). Nevertheless, Fouet and colleagues recently reported the isolation of an attenuated variant of the M1T1 serotype possessing the CovSY39H mutation. While the wild-type (WT) S119 was isolated from blood and caused NF and STSS, the variant S126 mutant was isolated from the pharynx of a household contact, showing no symptoms (35). The authors demonstrated that CovSY39H poorly responded to Mg^2+^ or LL-37, suggesting that this mutation compromises CovS phosphatase activity (35).

We combined RNA sequencing (RNA-seq), quantitative CovR phosphorylation assays, measurements of ScpC (also known as SpyCEP) activity, and a murine model of human GAS NF. We demonstrated that the CovSY39H mutant exhibits highly impaired phosphatase activity, yet retains its kinase and phosphotransferase activities. Moreover, we reversed the CovSY39H attenuated phenotype to that of the WT S119 by ectopically expressing the cytosolic domain of CovS in the S126*cov*S-3’ strain.

## RESULTS

### Characterizing gene transcription of the CovSY39H variant (strain S126) and evaluating its virulence in a subcutaneous murine model

We compared the growth of S126 and its parental strain, S119, in a chemically defined medium (CDM) supplemented with or without asparagine (Asn). Both strains exhibited enhanced growth kinetics in CDM supplemented with Asn, and both reached similar growth levels regardless of the presence or absence of Asn (Fig. 1a). However, the two strains exhibited significant differences in their RNA-seq profiles when grown to an optical density at 600 nm (OD_600_) of 0.7 in CDM with or without Asn (the results for all the detected genes are available under GEO accession numbers GSE296586 and GSE268517). For S126, out of the total number of detected genes (n=1784), 33.24% (n=593) were affected; among these, 13.97% (n=344) were upregulated, and 19.28% (n=249) were downregulated by at least 2-fold in Asn-supplemented versus non-supplemented conditions. For S119, out of the total number of detected genes (n = 1784), 27.13% (n = 484) were affected; of these genes, 13.84% (n = 247) were upregulated, and 13.28% (n = 237) were downregulated under similar conditions. The comparative transcriptome of S119 and S126 revealed that 21.63% (n=386) were upregulated and 17.04% (n=304) were downregulated by at least 2-fold in S119 when the strains were grown in an Asn-non-supplemented condition. In contrast, 6.22% (n=111) were upregulated and 5.49% (n=98) were downregulated by at least 2-fold in S119 when grown in Asn-supplemented conditions (GEO accession numbers GSE296586 and GSE268517).

**Figure 1.**
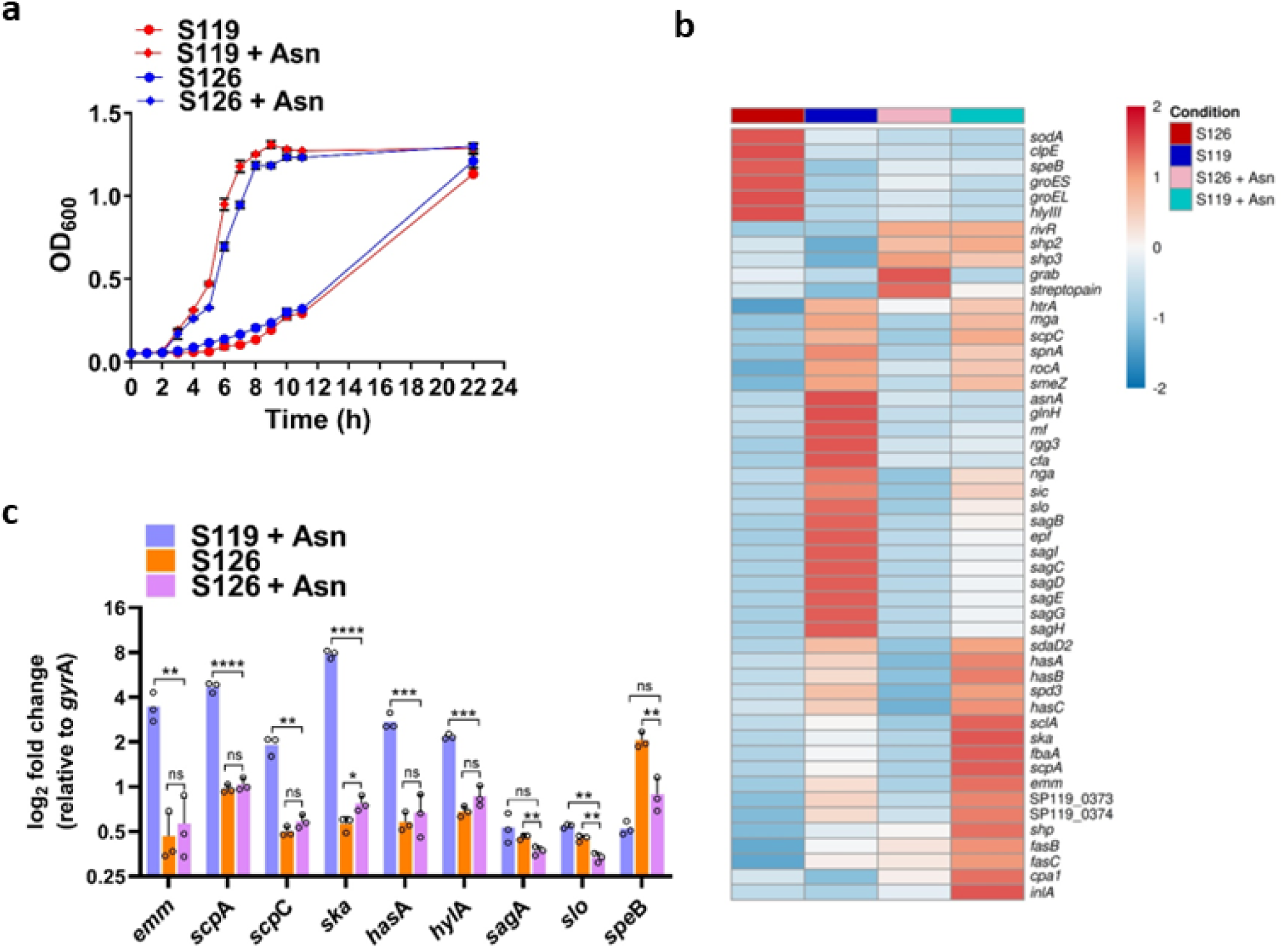
Comparative transcription analyses of GAS strains S119 (wild type) and S126 (CovSY39H variant). (a) The growth of the GAS strain S119 and S126 was determined in CDM in the absence or presence of Asn (10 µg/ml). (b) The heatmap displays differential gene expression patterns derived from RNA-seq data. Data illustrate the differential expression of selected genes in S119 and S126 when grown in CDM in the absence or presence of Asn (10 µg/ ml). (c) Quantitative real-time PCR (qRT-PCR) determinations of a few genes used in the RNA-seq analysis. In all qRT-PCR data, transcript abundance for each gene was normalized to that of *gyr*A in each sample, and fold changes were calculated in comparison with the normalized transcript abundance of the S119 grown without Asn. Three (a, c) and four (b) biological replicates were used. The values shown represent the means ± SD. Statistical analysis was performed using an unpaired two-tailed t-test (c).

Principal component analysis (PCA) was performed on RNA-seq data to examine variation across different samples. The analysis revealed that the biological replicates for each strain and condition clustered closely together, indicating high reproducibility and clear separation between experimental groups (Supplementary Fig. 1a). Volcano plots showed differential gene expression from RNA-seq comparisons. Each dot represents a gene with an adjusted P-value < 0.05, showing the fold change in gene expression between S119 and S126 grown in CDM in the absence or presence of Asn (10 µg/ml). The top three most highly expressed genes in S119 vs. S126 for each condition are shown in volcano plots (Supplementary Fig. 1b).

The heatmaps and qPCR data comparing the gene transcription of specific genes in S126 and S119, grown in CDM in the absence and presence of Asn, are shown in Fig. 1b. Without Asn, genes involved in stress survival, including *clp*E, *sod*A, *gro*EL, *gro*ES, and *hly*III were strongly upregulated in the S126 strain (Fig. 1b). The same pattern was observed for *spe*B, encoding a cysteine protease known as a virulence factor of GAS NF pathogenesis (36). In contrast, the WT S119 did not upregulate these genes under the same growth conditions (Fig. 1b). The transcription of the indicated genes was downregulated in S126 in the presence of Asn. However, the transcription of *riv*R, *SHP*2, and *SHP*3 was upregulated in both strains in the presence of Asn (Fig. 1b). A similar pattern was observed for the *fas* genes and *cpa*I (Fig. 1b), suggesting their regulation is independent of the CovS mutation Y39H. The Asn-mediated downregulation of the genes whose products participate in scavenging of Asn from the host and Asn metabolism, including SLO and SLS (*sag* genes) and asparagine synthetase (*asn*A), was significantly downregulated by the presence of Asn in S119 (Fig. 1b, c), as previously reported by us (37). Their transcription was also reduced in the S126 CovSY39H mutant, although their level of transcription was much lower except for *spe*B (Fig. 1c). Finally, genes involved in virulence, such as *has* genes, *ska*, *scl*A, *scp*A, and *emm,* were hardly affected by Asn in S126 but were strongly upregulated in S119 in the presence of Asn (Fig. 1b, c), as previously observed (37).

Since S126 virulence was assessed in the intravenous murine model (38), which may not accurately reflect the pathogenesis of invasive soft-tissue GAS infections, such as NF (39), we reassessed the virulence of S126 compared to that of S119 in a non-lethal murine model of human GAS NF (39). We enumerated GAS colony-forming units (CFUs) in soft tissue and spleen 3, 6, 9, and 11 days after subcutaneous inoculation (Fig. 2a). We also measured the lesion area and pictured the lesions (Fig. 2b and Fig. 2c). In addition, we determined the spleen weights (Supplementary Fig. 2a). All the results confirmed that S126 is severely attenuated compared to S119 in the murine model of human GAS NF. To determine whether the CovSY39H mutation is solely responsible for the attenuated phenotype, we swapped the wild-type CovS with the mutated CovSY39H, resulting in the strain S119*cov*S_126_. Indeed, the S119*cov*S_126_ strain was attenuated, generating significantly lower CFU in biopsies taken at 6, 9, and 11 days after subcutaneous inoculation (Supplementary Fig. 2b) and producing reduced lesion sizes (Supplementary Fig. 2c) compared to S119. Finally, we showed that the host immune responses of IL-1β, MIP-2, and IL-6 were also significantly lower in mice challenged with the variant S126 compared to those challenged with the WT strain S119 (supplementary Figs. 2d-f). In summary, the results taken together indicate that S126 is attenuated in a mouse model of human NF due to the single CovSY39H mutation.

**Figure 2.**
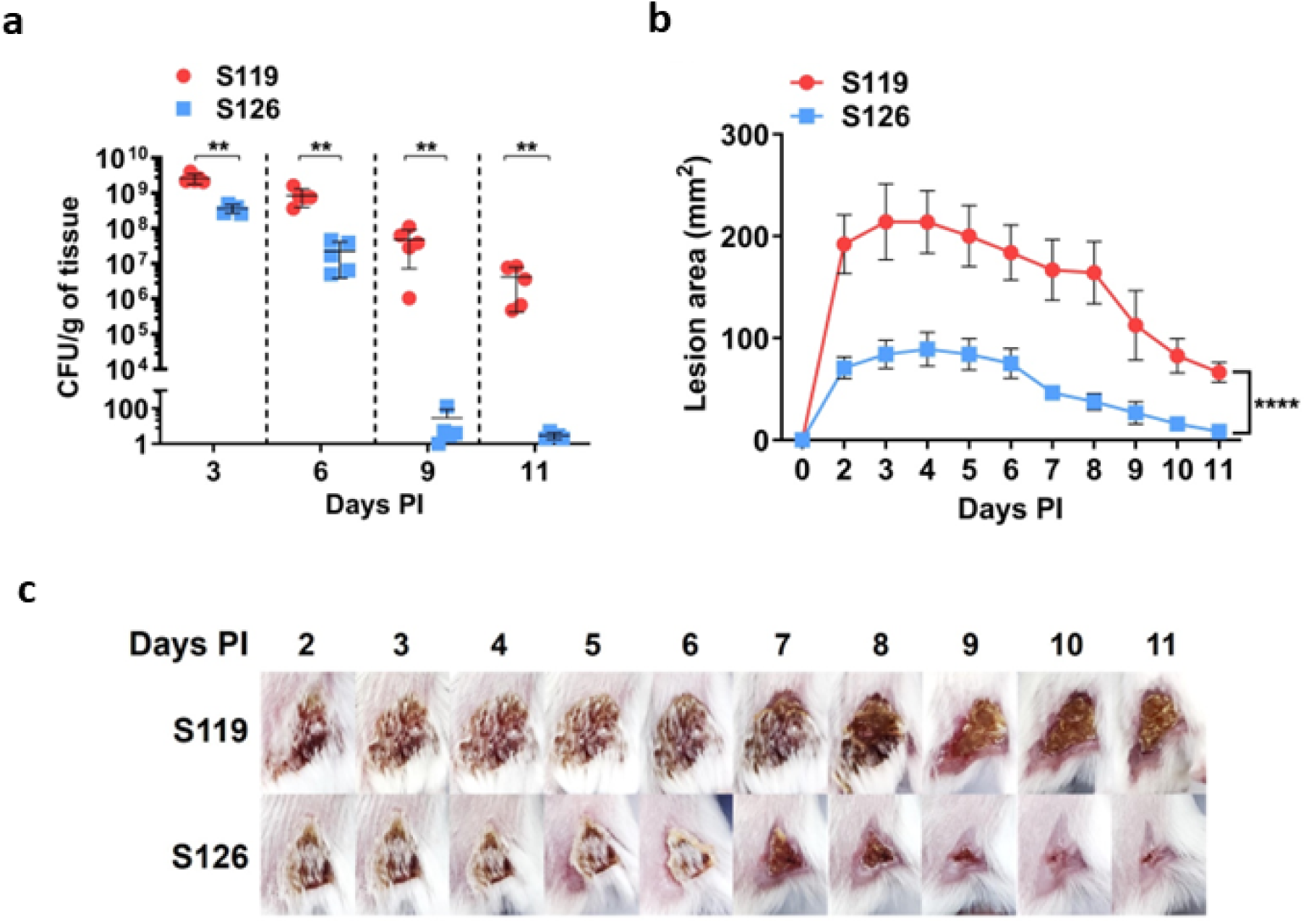
The CovSY39H mutant is attenuated in the GAS sublethal murine model of human NF. (a) BALB/c mice were injected with a sub-lethal dose of GAS through the subcutaneous (SC) route. CFU counts per gram of soft tissue derived from mice infected with S119 and S126 (CovSY39H) were enumerated and compared at different time points. (b) Lesion areas of mice infected with S119 and S126 were compared at different time points post-infection. (c) Mice were injected subcutaneously, and representative images of lesion progression are shown and compared at the indicated time points after infection with S119 and S126. Five mice were used per group per data point (a-c). The values shown represent the means ± SD. Statistical analysis was performed using the Mann–Whitney U test (A) and two-way ANOVA (b).

### The CovR of the *cov*SY39H variant is held in a phosphorylated state

CovS possesses three distinct activities. It phosphorylates CovR by histidine kinase (HK), transfers the phosphoryl group from histidine (His) 280 of CovS to aspartate (Asp) 53 of CovR, and dephosphorylates CovR∼P through its phosphatase activity (please see the diagram in supplementary Fig. 4a). Thus, the degree of phosphorylation at Asp53 of CovR reflects a steady-state level of this three-reaction system (17, 32). We measured phosphorylated CovR levels (CovR∼P/total CovR) in the CovSY39H variant strain S126 compared to its parental strain S119 by Phos-Tag gel electrophoresis technology (40). The comparison of representative Western blot demonstrates that the CovR of S119 was phosphorylated when grown in CDM, and its phosphorylation diminished when grown in CDM supplemented with Asn or Asn plus LL-37. However, no significant dephosphorylation was observed for CovR of S126 under the same conditions (Fig. 3a, b). These data demonstrate that CovSY39H exhibits histidine kinase (HK) and phosphotransferase activities but displays a significantly diminished phosphatase activity, as previously hypothesized (35).

**Figure 3.**
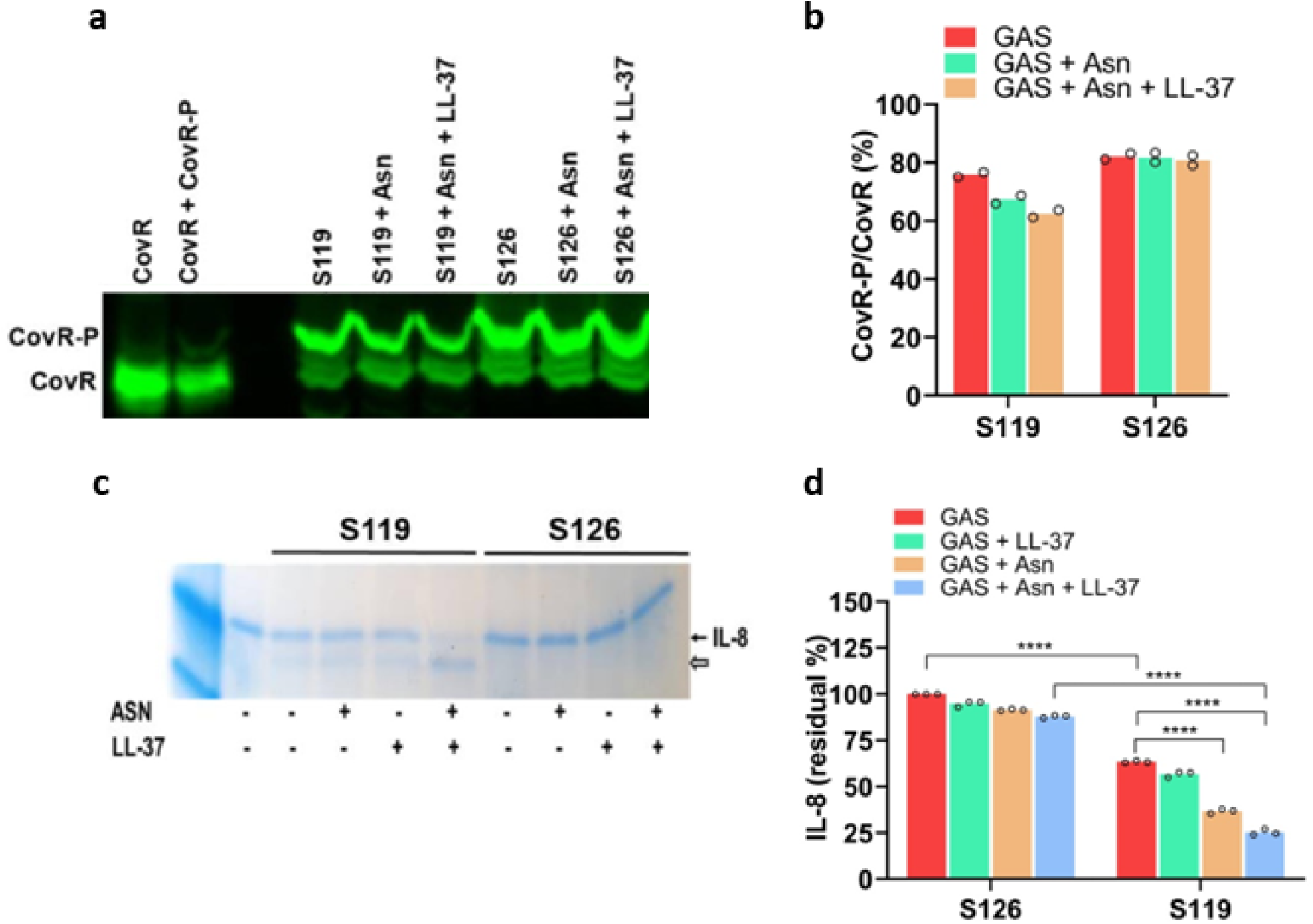
The CovR of S126 (CovSY39H variant) is held in a phosphorylated state. (a) CovR of the S126 strain is highly phosphorylated in comparison to the CovR of the S119 strain when grown in CDM without or with Asn or LL-37. Cell lysates were separated by Phos-Tag SDS-PAGE, with unphosphorylated (lane 1, from left) and phosphorylated recombinant CovR protein (lane 2, from left). CovR expression was detected using an anti-CovR antibody and visualized using a fluorescently labeled secondary antibody. (b) The percentages phosphorylated CovR (CovR-P) relative to total CovR were calculated using ImageJ software. (c) S126 does not display significant ScpC activity in culture media. Culture media of S119 and S126 were collected after growth in the absence or presence of Asn or/and LL-37 and then subjected to ScpC-mediated cleavage of recombinant human IL-8, followed by SDS-PAGE on Tris-tricine gels. The gels were visualized using Coomassie blue staining. (d) The determinations of IL-8 residual content in the supernatants of the indicated strains cultured without or with Asn or/and LL-37 were quantified by ELISA. Two biological replicates (a, b) and three (d) were used. The values shown represent the means ± SD. Statistical analysis was performed using an unpaired two-tailed t-test (d).

The reduction in CovR phosphorylation (CovR∼P/total CovR) in S119 was significant compared to that of the CovR in the S126 variant, which barely but insignificantly decreased in the presence of Asn + LL-37. However, the differences in phosphorylation levels ranged only from approximately 78% to 64% (Fig. 3b). We tested whether these changes could be amplified at the level of virulence factor production. To this end, we measured the activity of GAS CXC-chemokine protease, ScpC (also known as SpyCEP), which is directly regulated by CovR/S (17, 18). First, we visualized the cleavage of IL-8 by exposing the purified chemokine to culture media of S119 or S126 grown in CDM in the absence or presence of Asn, LL-37, or both. From the SDS-PAGE shown in Fig. 3c, it is apparent that the culture media of the S119 contained ScpC activity, which was most evident when the bacteria were grown in the presence of both Asn and LL-37. Next, we quantified ScpC activity by ELISA. Adding Asn and LL-37 to CDM generated the highest ScpC activity for strain S119. There was a slight increase in ScpC activity for strain S126 under the same conditions, but it was insignificant (Fig. 3d). C5a degradation by ScpA was also apparent in the culture media of S119 when grown in the presence of Asn and LL-37 (Supplementary Fig. 3a).

### Expressing the cytosolic domain of CovS suppresses the phosphatase inactivation of the CovSY39H variant

We attempted to restore S126 CovS phosphatase activity by ectopically expressing the cytosolic domain of CovS in strain S126. This notion was based on **(a)** previous findings showing that the cytosolic domains of structurally related kinase/phosphatase sensors express phosphatase activity in the presence of ADP (30, 31), **(b)** our recent findings showing that GAS grown in CDM supplemented with Asn produces an excess of intracellular ADP over ATP (37). To this end, we cloned the cytosolic domain of CovS, encoding 282 amino acids (aa) (aa 218 to aa 500), into a shuttle plasmid, pKSM411, harboring the *emm1* promoter (41) and transformed S126 with the resulting plasmid, yielding the strain S126pKSM-*cov*S-3’ (see Supplementary Fig. 4b for the corresponding illustration). First, we examined the transcription level of several genes of S126pKSM-*cov*S-3’, which were activated in S119 and not in S126 when grown in CDM supplemented with Asn (Fig. 1b). We found that these genes were strongly activated in S126pKSM-*cov*S-3’ (Fig. 4a). As anticipated, the expressed cytosolic CovS fragment (aa 218 to aa 500) produced phosphatase activity (Fig. 4b), which dephosphorylated the CovR of S126pKSM-*cov*S-3’, causing the increased transcription of the indicated virulence factors (Fig. 4a). The dephosphorylation of CovR decreased from about 78% to 58% (Fig. 4c), which was larger compared to dephosphorylation of the strain S119 grown under the same conditions (Fig. 3a, b). These results support the notion that the functional activity of the phosphatase expressed from the plasmid pKSM-*cov*S-3’ was greater than that expressed from the chromosome of S119. To substantiate this notion, we measured ScpC activity of S126pKSM-*cov*S-3’. Indeed, we found that ScpC activity was apparent even when grown in CDM alone without any supplementation. Additionally, it was strongly upregulated by supplementing CDM with Asn compared to S119 under all growth conditions tested (Fig. 5a). Moreover, when the *cov*S-3’ fragment was expressed in the S119 background, yielding the S119pKSM-*cov*S-3’ strain. Its ScpC activity was substantially higher than that of S119 under all growth conditions tested. This data supported the notion that the phosphatase activity expressed from the plasmid acts in trans to dephosphorylate CovR, which is expressed from the chromosome. We also mutated the active site of CovS (CovS-H280V), which obliterates its kinase/phosphatase activity (40). The resulting strain, S126pKSM-*cov*S-3’_H280V_, did not produce any ScpC activity under the tested growth conditions (Fig. 5b). Furthermore, it acted as a dominant negative factor in the S119 background since the strain S119pKSM-*cov*S-3’_H280V_ lacked any ScpC activity as well (Fig. 5b).

**Figure 4.**
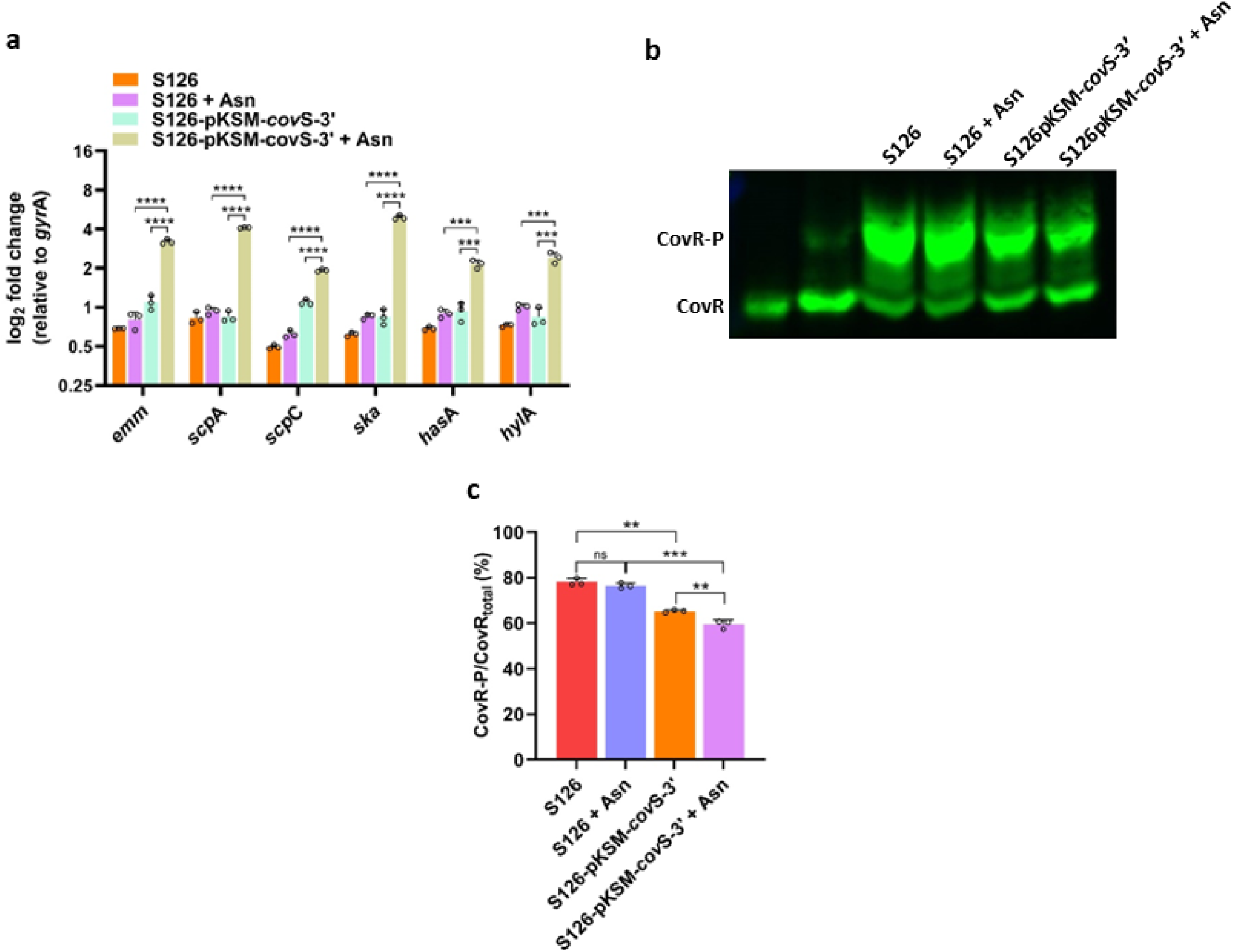
Expression of the cytosolic domain of CovS suppresses the phosphatase inactivation of the CovSY39H variant. (a) The cytosolic domain of CovS encoding 282 amino acids (aa) (aa 218 to aa 500) was expressed from the *emm*1 promoter in S126 background, yielding the strain S126pKSM-*cov*S-3’. qRT-PCR was performed to assess the transcription of the corresponding virulence genes. In all qRT-PCR data, transcript abundance for each gene was normalized to that of *gyr*A in each sample, and fold change was calculated in comparison with the normalized transcript abundance of the S119 grown without Asn. (b) CovR was significantly dephosphorylated in S126pKSM-*cov*S-3’ in comparison to S126 when grown in CDM without or with Asn. Cell lysates were separated by Phos-Tag SDS-PAGE, with unphosphorylated (lane 1, from left) and phosphorylated recombinant CovR protein (lane 2, from left). CovR expression was detected using an anti-CovR antibody and visualized using a fluorescently labeled secondary antibody. (c) The percentages of CovR-P of total CovR protein were calculated using ImageJ. For (a-c), three biological replicates were used. The values shown represent the means ± SD (a, c). Statistical analysis was performed using an unpaired two-tailed t-test (a, c).

**Figure 5.**
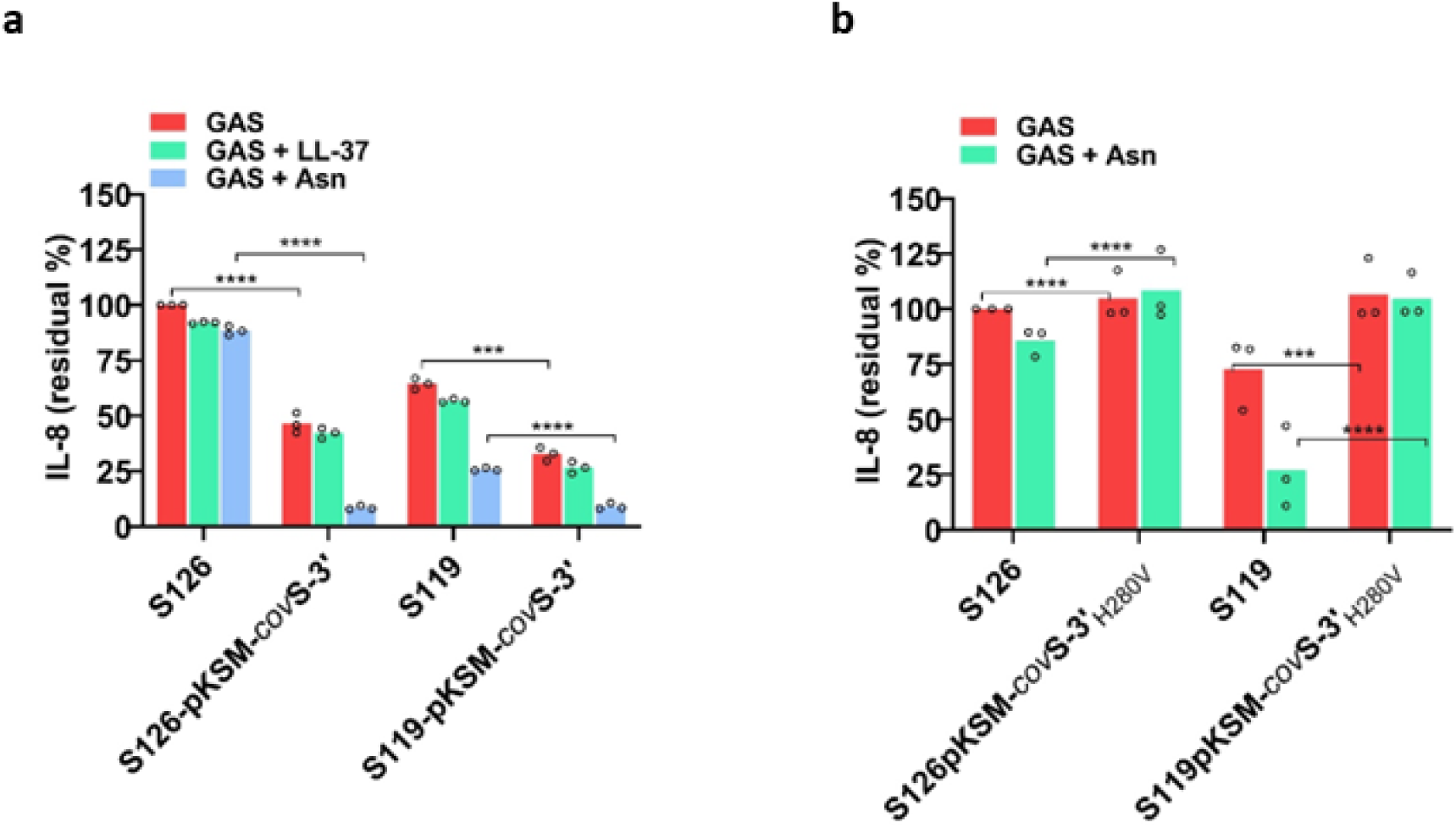
The cytosolic domain of CovS expressed from the plasmid acts in trans on CovR expressed from the chromosome. (a) CovS encoding the cytosolic domain (aa 218 to aa 500) was expressed from a plasmid in S126 and S119 backgrounds (strains S126pKSM-*cov*S-3’ and S119pKSM-*cov*S-3’, respectively). (b) The same plasmid harboring a mutation in the active site (H280V) was expressed in the S126 and S119 backgrounds (strains S126pKSM-*cov*S-3’_H280V_ and S119pKSM-*cov*S-3’_H280V_). The determinations of IL-8 residual content in the supernatants of the indicated strains cultured without or with Asn or LL-37 were measured by ELISA. Three biological replicates (a, b) were used. The values shown represent the means ± SD. Statistical analysis was performed using an unpaired two-tailed t-test (a, b).

Additionally, we assessed the virulence of S126pKSM-*cov*S-3’ using the mouse non-lethal model of human NF (39). We enumerated CFU and estimated the lesion sizes of mice injected with strains S119, S126, and S126pKSM-*cov*S3. As expected from Fig. 2, biopsies taken after 2, 6, and 9 days from mice challenged with S126 contained much lower CFU compared to mice challenged with S119 on days 6 and 9 after challenge (Fig. 6a). The CFU of mice challenged with S126pKSM-*cov*S-3’ were similar to those of mice challenged with S119 (Fig. 6a). The same was true for the lesion sizes (Fig. 6b). All colonies isolated from mice challenged with S126pKSM-*cov*S-3’ harbored the plasmid pKSM-*cov*S-3’, probably because it endows the corresponding strain with a high survival advantage within the mouse soft tissue.

**Figure 6.**
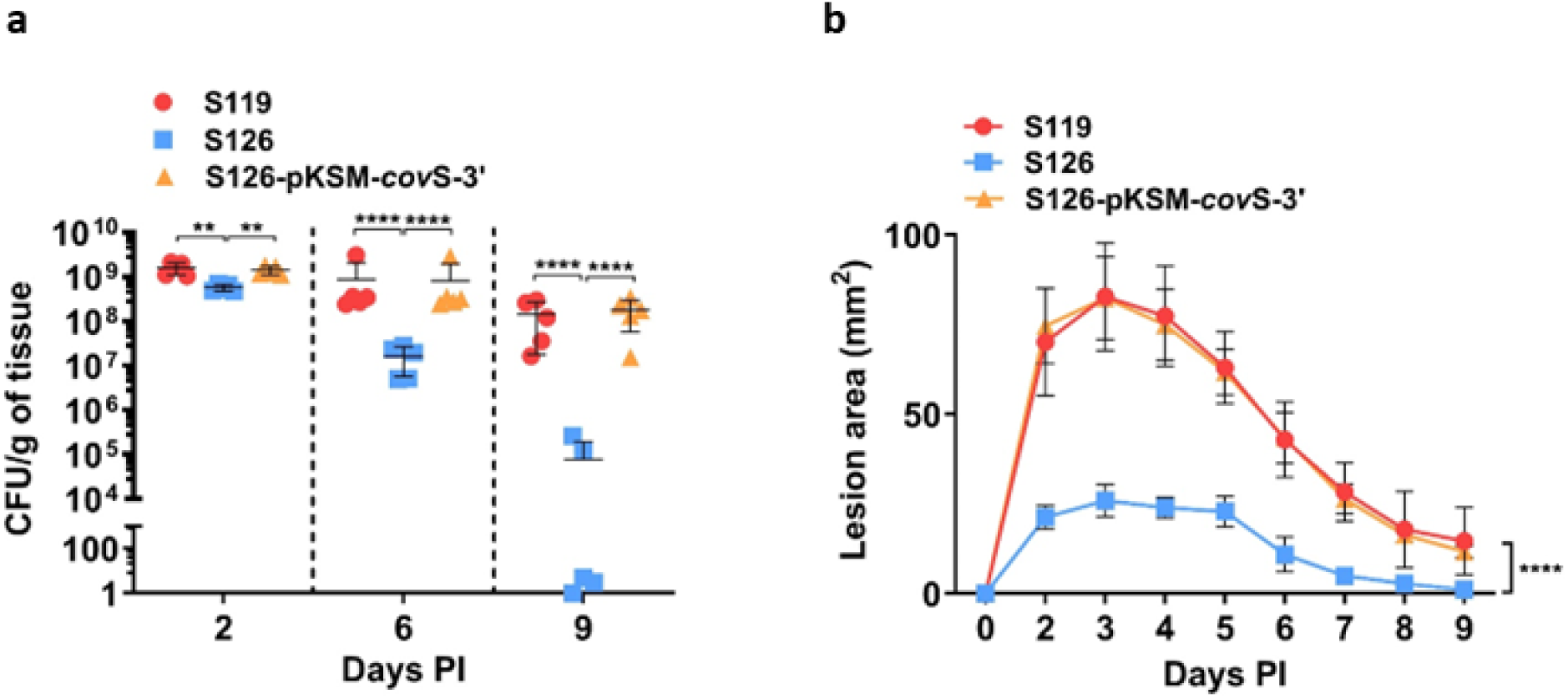
S126pKSM-*cov*S-3’ was as virulent as S119 in a sublethal murine model of human GAS NF. (a) BALB/c mice were injected with a sub-lethal dose of GAS through the subcutaneous (SC) route. CFU counts per gram of soft tissue derived from mice infected with S119, S126 (CovSY39H), and S126-p*KSM*-*cov*S-3’ were enumerated and compared at different time points. (b) Lesion areas of mice infected with S119, S126 (CovSY39H), and S126-p*KSM*-*cov*S-3’ were compared at different time points post-infection. Five mice were used per group per data point (a, b). The values shown represent the means ± SD. Statistical analysis was performed using the Mann–Whitney U test (A) and two-way ANOVA (B).

## DISCUSSION

A colonizer M1T1 GAS variant that harbors a mutation in CovS-Y39H (strain S126) was isolated in France and found to be attenuated compared to its parental wild-type strain S119, causing NF and STSS. The authors hypothesized that the mutation causes a loss of CovS phosphatase activity (35).

Here, we performed a genome-wide RNA-seq comparison of S126 and S119 grown in CDM to an optical density (OD) of 0.7 in the presence and absence of Asn. We found that CovR/S-regulated virulence genes were almost unresponsive to Asn in S126, whereas in S119, the transcription of these genes was strongly upregulated, as recently reported (37). We also discovered that S126 exclusively upregulated the transcription of the stress response genes *clp*E, *sod*A, *gro*EL, *gro*ES, and *hly*III in the absence of Asn. Since it was reported that CovR/S contributes to stress tolerance (42), the CovSY39H variant (S126) probably faces higher stress levels than the parental S119 strain when grown in CDM without Asn. Interestingly, in the presence of Asn, the transcription of the indicated genes was downregulated in S126, suggesting that Asn uptake by GAS may also alleviate stress independently of CovR/S signaling.

In GAS, the differential gene expression and *in vivo* virulence are primarily regulated by the CovR/S two-component system (TCS), which acts mainly through gene repression (16, 24) and enables GAS to respond to host cues (15, 17, 18, 20, 33). Indeed, a direct interaction between CovS and LL-37 has been reported to alter gene transcription by dephosphorylating CovR, thereby increasing CovS phosphatase activity (17, 22, 32). Recently, we demonstrated that the presence of Asn during the growth of GAS in CDM reduces CovR phosphorylation, due to the formation of a surplus of ADP over ATP caused by the uptake and synthesis of Asn. We suggested that an excess of ADP stimulates the phosphatase activity of CovS (37), as observed in structurally related kinase/phosphatase sensors (27–31). Here, we verified this notion directly.

To achieve this, we utilized Phos-tag technology, which enables the quantitative monitoring of phosphotransfer reactions from HK to CovR despite the instability of phosphorylated histidine and aspartate (43, 44). We demonstrated that in S126, both the HK and phosphotransfer activities of CovS were intact, as CovR remained phosphorylated in the presence of Asn, LL-37, or both compounds. Thus, as predicted, the phosphatase activity of CovS-Y39H is impaired, and the S126 mutant is attenuated. The mechanism by which the mutation in amino acid His-39, located at the extracellular site very close to the membrane surface (see Supplementary Fig. 4a), causes inactivation of the phosphatase activity in S126 is currently unknown. In the PhoP/PhoQ TCS, which resembles the CovR/S structure, a scissor-like movement in PhoQ has been suggested to be responsible for signal transduction to PhoP, leading to the dynamic rearrangement of the transmembrane helices (45, 46). Perhaps such movement is restricted in the S126 mutant, allowing HK phosphorylation and phosphotransfer but inhibiting phosphatase reaction, which requires a more substantial movement (45, 46). A comparative structural analysis of CovS from S119 and S126 may answer this question.

The phosphatase activity of the cytosolic portion of EnvZ, homologous to CovS, is strongly stimulated by ADP)31 ,30(; therefore, we decided to ectopically express the cytosolic portion of CovS in an attempt to reverse the phenotype of S126 to that of S119. We hoped for a positive outcome because an excess of ADP is formed over ATP in GAS grown in CDM supplemented by Asn (37). Indeed, the S126*cov*S-3’ phenotype fully reverted to that of S119 in terms of CovR/S-mediated gene regulation and virulence factor production. Most importantly, it also reverted *in vivo*, shifting from an attenuated strain into a fully virulent one, demonstrating that Asn-mediated GAS virulence regulation occurs *ex vivo* and *in vivo* in an animal model mimicking human GAS NF. Our study provides a novel mechanistic tool that enables the manipulation of CovS phosphatase activity without affecting its kinase activity.

## MATERIALS AND METHODS

### Bacterial culture

The GAS strains used in this study are represented in Supplementary Material Table 1. The methodology and primers used for constructing all mutants are described in Supplementary Materials Table 2. GAS was cultured overnight without shaking in Todd-Hewitt broth supplemented with 0.2% yeast extract (THY) in sealed tubes at 37°C. When necessary, antibiotics were added at the final concentrations of 250 µg/ml for kanamycin (Km) or 50 µg/ml for spectinomycin (Spec). The following morning, overnight cultures were diluted 1:20 and grown in THY medium with the appropriate antibiotic, as needed, to an early log phase (OD_600_ of 0.3). They were then washed and resuspended in chemically defined medium (CDM) designed by van de Rijn and Kessler (47). The growth rates of different bacterial strains were determined in CDM supplemented with or without Asn at concentrations of 0 or 10 µg/ml. 1 mL of freshly prepared CDM (in the absence or presence of Asn and an appropriate antibiotic) was added to each well of a 24-well plate, and the plate was seeded with bacteria and incubated at 37°C in a 5% CO_2_ atmosphere. The absorbance was measured at an optical density (OD) of _600_ at regular time intervals. This method of culturing GAS in CDM was followed for all experiments.

### Plasmid construction

For swapping the CovS of S119 with that of S126 in the S119 background, we constructed the pJRS*cov*S_126_ plasmid. A 717 bp fragment containing 107 bp of the end of *covR* and 611 bp of the beginning of *cov*S was amplified with the primers *cov*R/S-p3-F and *cov*R/S-p9-R and using S126 genomic DNA as a template. The PCR product was AT cloned into pGEM-T® Easy (NEB, USA), yielding the plasmid pGEM*covS*_126_. Next, pGEM*covS*_126_ was digested with *NotI* and *PstI,* and the resulting *covS* fragment was ligated into pJRS233, which had been digested with the same enzymes. The plasmid pKSM-*cov*S-3’ was constructed using Gibson assembly. A 852 bp fragment of *cov*S-3’ (aa 218-end) was amplified from S119 genomic DNA using primers *cov*S 218-end -F and *cov*S 218-end -R (Supplementary Materials Table 2). A 6773 bp fragment of the plasmid pKSM411 (48), excluding the *gfp* gene, was amplified using the primers pKSM411-*cov*S-Gibson-F and pKSM411-*cov*S-Gibson-R (Supplementary material Table 2). The amplified PCR products were ligated using the Gibson assembly kit (NEB, USA). The resulting plasmid expressed 3’ *cov*S from the *emm*1 promoter. The pKSM-*cov*S-3’_H280V_ plasmid was constructed using the Gibson assembly method. A 3618 bp fragment containing a portion of the *cov*S 3’ region (amino acids 218 to the end) along with part of the pKSM backbone was amplified from the pKSM-*cov*S-3’ plasmid using primers SpecR-F and *cov*S 218-end-H280V-R (Supplementary material Table 2). Additionally, a 4017 bp fragment comprising the remaining part of the *cov*S 3’ region and the remaining part of the pKSM plasmid was amplified using primers SpecR-R and *cov*S 218-end-H280V-F (Supplementary Materials Table 2). The two PCR fragments were then assembled using the Gibson assembly kit (NEB, USA). The final plasmid expresses the 3’ *cov*S region carrying a histidine-to-valine point mutation at position 280, driven by the *emm*1 promoter.

### Bacterial strain construction

S119*cov*S_S126_ (119 harboring CovS of S126 ) was constructed by transforming pJRS*cov*S126 into S119Δ*cov*S (49) as described before (50). S119pKSM-*cov*S-3’, S126pKSM-*cov*S-3’, S119pKSM-*cov*S-3’_H280V_, and S126pKSM-*cov*S-3’_H280V_ strains were obtained by introducing the pKSM-*cov*S-3’ or pKSM-*cov*S-3’_H280V_ plasmids into GAS S119 and S126, respectively, through electroporation. Bacterial colonies containing the plasmid were selected by growth at 37°C on THY agar plates supplemented with either Spec or Km.

### Extraction of RNA from GAS and qRT-PCR determinations

The total RNA of GAS was isolated using the phenol-ethanol extraction method, and purification was conducted using the Direct-zol RNA miniprep kit (Zymo Research). The RNA concentration and purity were evaluated using a NanoDrop One spectrophotometer (Thermo Scientific). RNA was treated with RQ1 DNase (Promega) according to the manufacturer’s instructions to avoid genomic DNA contamination. According to the manufacturer’s instructions, M-MLV Reverse Transcriptase (Promega) was used for reverse transcription. For real-time PCR, the cDNA was diluted, and quantitative PCR was performed on a Mic qPCR Cycler (Bio-Molecular Systems) using 2X Tamix Fast SyGreen Mix Hi-ROX (Tamar Laboratory Supplies Ltd.). The primers used are listed in Supplementary Materials Table 3. The expression levels of each target gene were normalized to *gyr*A and analyzed using the 2-ΔΔC_T_ method (51).

### RNA-Seq and data analysis

Total RNA was extracted from GAS using the phenol-ethanol extraction method, and purification was conducted using the Direct-zol RNA miniprep kit (Zymo Research). Ribosomal RNA was removed using the Pan-Bacteria (RNA-Seq) riboPool^TM^ kit (siTOOLs BioTech, Germany). Sample quality was assessed using a 2100 Bioanalyzer (Agilent), and sample quantity was determined using a NanoDrop 8000 spectrophotometer (Thermo Scientific). According to the manufacturer’s recommendations, the RNA-seq directional libraries were generated using the KAPA Stranded mRNA-Seq Kit (Kapa Biosystems, KK8421). A 76 bp single-read DNA sequencing was performed using the Illumina Nextseq500 platform (Core Research Facility, Faculty of Medicine, The Hebrew University of Jerusalem, Israel). Data were generated in the standard Sanger FastQ format, and raw reads have been deposited under BioProject accession numbers GSE296586 and GSE268517 with the Sequence Read Archive (SRA) at the National Center for Biotechnology Information (NCBI). NextSeq basecalls files were converted to fastq files using the bcl2fastq program with default parameters (without trimming or filtering applied at this stage). After ensuring quality, the processed reads were aligned to the reference genome of *Streptococcus pyogenes* strain S119 (GCA_900608505.1, GenBank).

### ScpC and ScpA cleavage assay

GAS was cultured in CDM in the absence or presence of Asn (10 µg/ml) or 300 nM LL-37 in 24-well plates at 37°C in 5% CO_2_ atmosphere, and samples were collected at late log phase (OD_600_ of 0.7). The cell-free supernatants were incubated with an equal volume of 1 mg/ml purified recombinant human IL-8 (R&D Systems, USA) and human complement component C5a (R&D Systems, USA) at 37 °C for 2 hours. The samples were heated at 100°C for 5 min with 4x Tricine loading buffer (0.4 M Tris HCl pH 6.8, 80% glycerol, 4% SDS, and 0.08% Coomassie blue) to stop the reaction. The proteins were resolved on precast 16.5% Mini-PROTEAN® Tris-Tricine gels using the Mini-PROTEAN gel apparatus (Bio-Rad). The samples were run at a low constant voltage of 4°C for 6 to 8 hours in Tris-Tricine SDS Running Buffer (Bio-Rad, USA). Peptides were detected using InstantBlue (Expedeon Inc.). The gels were destained with distilled water until the bands were visible.

### ELISA-based assessment of ScpC expression by performing IL-8 degradation assays

The samples from GAS culture were collected as mentioned above, and the cell-free supernatants were incubated with 1 ng/ml of recombinant human IL-8 (R&D Systems, USA) at 37°C for 2 h. The samples were heated at 100°C for 2 min; the reaction was stopped. IL-8 cleavage was represented as the residual amount of IL-8 in the samples, estimated using the Human IL-8 Quantikine ELISA kit (R&D Systems, USA). The IL-8 residual content in all supernatants was normalized to that of the GAS S126 strain without Asn.

### Protein extraction and Phos-tag Western blotting

Bacteria were cultured in CDM without or with Asn (10 µg/ml) until the OD_600_ reached 0.6 to 0.7. Subsequent steps were performed at 4°C. Bacterial cultures were collected, centrifuged, and the pellets were washed with ice-cold sterile PBS. For protein extraction, the pellet was suspended in ice-cold lysis buffer [20 mM Tris-HCl, pH 8, with 10 U mutanolysin (Sigma Aldrich), Complete™ EDTA-free Protease Inhibitor Cocktail (Roche Molecular Diagnostics, USA), and PhosSTOP™ (Roche Molecular Diagnostics, USA) in PBS]. The bacterial cells were homogenized by using MagNA Lyser (Roche Diagnostics). The lysates were centrifuged to collect the supernatants, and the total protein concentration was quantified using the Bradford assay. Protein samples were mixed with 4x Laemmli Sample Buffer (900 µl of Laemmli Sample Buffer + 100 µl of ß-mercaptoethanol) and resolved in Phos-tag SuperSep Phos-tag Gels (Wako, Japan). Recombinant CovR was purified and phosphorylated *in vitro* as described (52) and served as a control. The gel was treated three times with blotting buffer [10x SDS PAGE buffer 50mL (5 of 1X) + MetOH 200 mL (2) + adjusted to a total volume of 1 liter (milliQ)] supplemented with 10 mM EDTA to remove all Zn^2+^ from the gel. The blotting was performed using the Trans-Blot Turbo Transfer Pack (Bio-Rad). The membrane was blocked overnight at 4°C in PBST with 5% skim milk. The membrane was washed and then placed in a suspension of rabbit anti-CovR antibody in PBS-T (1:5000) for 1 hour at room temperature. The membrane was washed and treated with a fluorescently labeled Goat Anti-Rabbit IgG StarBright Blue 700 antibody (1:5000) in 10 mL of PBST buffer with 1% skim milk at room temperature for 2 hours. Finally, the membrane was washed, and the fluorescently labeled proteins were visualized at a wavelength of 470 nm using the Biorad GelDoc system. The relative percentage of phosphorylated CovR was calculated using ImageJ software (open source, developed by NIH, USA).

### Animals

3 to 4-week-old female BALB/c OlaHsd mice weighing 10-12 g were obtained from ENVIGO RMS (Israel Ltd.). Following the Hebrew University of Jerusalem’s ethical guidelines, all procedures were performed in accordance with the humane handling, care, and treatment of research animals (Protocol number MD-22-17143-5). Mice were kept in disposable cages supplemented with enrichment, and regular sterile food, water, and air were supplied separately to each cage. All cages were placed in specific pathogen-free (SPF) conditions during the experiment with controlled environmental conditions. The mice were left to acclimate for 3 days, after which the treatment groups were randomized, and the littermates were evenly distributed among the cages. Identification markings and shaving of the dorsal flanks of already weighed mice were performed, and the mice were then infected. Following infection, mice were fed wet food twice daily and monitored for parameters such as body weight, activity level, fur quality, and eye appearance. According to the guidelines of the Institutional Animal Care Units at the Hebrew University’s School of Medicine, based on the above parameters, a scoring method was implemented to determine humane endpoints, at which mice were euthanized in accordance with ethically approved procedures.

### Sublethal murine model of human GAS soft tissue infection

3- to 4-week-old female BALB/c OlaHsd mice weighing 10-12 g were obtained from ENVIGO RMS (Israel Ltd.). Following the Hebrew University of Jerusalem’s ethical guidelines, all procedures were performed in accordance with humane handling, care, and treatment of research animals (Protocol number MD-22-17143-5). The murine model of human soft-tissue infection was injected with a sub-lethal dose (5 × 10^7^ CFU) of GAS strains injected subcutaneously (SC) into the rear flank of mice, and CFU counts were determined in soft skin and spleen samples. At various time points, mice were euthanized by inhalation of isoflurane followed by cervical dislocation, and skin and spleen samples were collected. Skin tissue from the injection site was collected using a punch biopsy tool (Acu-Punch, Acuderm Inc.), and spleen samples were excised and transferred to 2 ml Eppendorf tubes containing 0.5 ml of sterile PBS. Tissues were homogenized, diluted, and plated on blood agar plates for all experiments, and CFUs were counted after overnight incubation at 37°C. CFU counts were normalized to the weight of the soft tissue. The CFU counts for S126-pKSM-*cov*S-3’ were matched with parallel plating on kanamycin (250 µg/µl) supplemented THY agar plates to check the stability of the extra-chromosomal plasmid. For determining the lesion area, dermonecrotic skin lesions were measured daily using a digital caliper (Bar Naor Ltd.). The lesion area was calculated with the formula: A = (π/2)(length)(width) (52).

### Cytokine measurements

Mice were infected subcutaneously with 5 × 10^7^ CFU of GAS in 100 μl of sterile PBS on the rear flank. After GAS infection, mice were euthanized at various time points, and soft tissue samples surrounding the lesion were harvested using a punch biopsy. The samples were minced with scissors and homogenized in lysis buffer containing 10 mM Tris-HCl (pH 7.8), supplemented with 1% NP-40, 150 mM NaCl, 40 mM EDTA, and Complete, Mini-EDTA-free protease inhibitor cocktail tablet. The tissue samples were vortexed for 1 hour at room temperature and centrifuged at 17,000g for 5 min. Supernatants containing chemokines and cytokines were collected and stored at −80°C. According to the manufacturer’s protocol, the amounts of chemokine MIP-2 and the pro-inflammatory cytokines IL-1β and 1L-6 were quantified using Quantikine enzyme-linked immunosorbent assay (ELISA) kits (R&D Systems). The amounts of chemokine and cytokine were normalized to the total protein content of the corresponding samples, as measured by the Bradford protein assay (Bio-Rad Laboratories).

### Statistical analysis

GraphPad Prism version 10 software was used to plot the results and perform statistical analysis. All values were represented as means ± standard deviation (SD). Data in bar graphs were analyzed using parametric unpaired two-tailed t-tests unless specified, and two-way ANOVA with Tukey post-tests and Mann–Whitney U test, where indicated. In all figures, P values were calculated to confirm significance (ns > 0.05, *P < 0.05, **P < 0.01, ***P < 0.001, ****P < 0.0001).

## ACKNOWLEDGMENTS

We thank the core facility of the Faculty of Medicine at Hebrew University for conducting the RNA-seq analysis. We are indebted to Dr. Abed Nasereddin (Core Research Facility, Faculty of Medicine, The Hebrew University of Jerusalem) and Dr. Yuval Nevo (Unit of Bioinformatics, Faculty of Medicine) for their valuable contributions. We thank Prof. Claire Poyart from Université Paris Cité, Institut Cochin, INSERM, U1016, CNRS, UMR8104, Paris, France, for providing the S119 and S126 GAS strains within the framework of the Complexity in Biology program, supported by the High Council for Scientific and Technological Cooperation between France and Israel. Finally, we thank Prof. Samuel A. Shelburne and Dr. Nicola Horstmann from the Department of Infectious Diseases, Infection Control, and Employee Health at The University of Texas MD Anderson Cancer Center, Houston, Texas, USA, for providing the necessary reagents and expertise for conducting the CovR phosphorylation assessment. This work was supported by the Israeli Ministry of Innovation, Science, and Technology (Grant number 0005663) and the Israel Science Foundation (ISF) (Grant number 1926/24) to E. H.

## SUPPLEMENTARY MATERIAL

Figures: S1-S4

Supplementary material Tables 1-3

